# Incorporating trait-based mechanisms, alternative interaction modes and forbidden links increases the minimum trait dimensionality of ecological networks

**DOI:** 10.1101/192260

**Authors:** Diogenis A. Kiziridis, Lynne Boddy, Daniel C. Eastwood, Chenggui Yuan, Mike S. Fowler

## Abstract

1. Species’ traits mediate ecological interaction outcomes and community structure. It is important, therefore, to identify the minimum number of traits required to characterise observed networks, i.e. the ‘minimum dimensionality’. Existing methods for estimating minimum dimensionality often lack three features commonly associated with a higher numbers of traits: a mechanistic description of observed interactions, alternative interaction modes (e.g. different feeding strategies such as active *vs* sit-and-wait feeding), and trait-mediated ‘forbidden links’. Omitting these features can lead to underestimation of the trait numbers involved, and therefore, minimum dimensionality. Here, we develop a ‘minimum mechanistic dimensionality’ measure, accounting for these three features.
2. The only input required by our method is the observed network of interaction outcomes. We then assume how traits are mechanistically involved in alternative interaction modes. These unidentified traits are contrasted using pairwise performance inequalities between the interacting species. E.g. if a predator feeds upon a prey species via a typical predation mode, in each step of the predation sequence the predator’s performance must be greater than the prey’s. We construct a system of inequalities from all observed outcomes, which we attempt to solve with mixed integer linear programming. The minimum number of traits required for a feasible system of inequalities provides our dimensionality estimate.
3. We applied our method to 658 published empirical ecological networks including animal dominance, predator–prey, primary consumption, pollination, parasitism and seed dispersal networks, to compare with minimum dimensionality estimates when the three focal features are missing. Minimum dimensionality was typically higher when including alternative interaction modes (54% of empirical networks), ‘forbidden interactions’ as trait-mediated interaction outcomes (92%), or a mechanistic perspective (81%), compared to network dimensionality estimates missing these features.
4. Our method can reduce the risk of omitting essential traits that are involved mechanistically, in different interaction modes, or in failure outcomes. More accurate estimates will allow us to parameterise models to generate theoretical networks with a more realistic structure at the interaction outcome level. Thus, we hope our method can improve predictions of community structure and structure-dependent dynamics.

## 1 INTRODUCTION

Communities are structured by different forces, including evolution, spatio-temporal distribution of species due to neutral, historical, dispersal, or habitat filtering processes, and species interactions (Bartomeus et al., 2016; Poisot, Stouffer, & Gravel, 2015; Vázquez, Chacoff, & Cagnolo, 2009). Traits of interacting species contribute to relative performance in the interaction, hence influencing the interaction outcomes, i.e. whether one species successfully exploits the other (Bartomeus et al., 2016; Pichler et al., 2019). For example, a nectarivory outcome can depend on the length of a nectarivore’s mouth part compared to the depth of the plant’s corolla tube (Maglianesi et al., 2014). Thus, the comparison of trait-mediated performance between the interacting exploiter–resource species underlies the interaction outcomes and, subsequently, community structure (Arnold, 1983). Here, we develop a method which leads backwards from the observed interaction outcomes to an estimate of the minimum number of traits that were involved in that type of interaction, which we term ‘minimum dimensionality’.

Knowing the minimum dimensionality for a set of interaction outcomes concentrates our efforts on investigating which traits underpin community structure (Eklöf et al., 2013). A set of interaction outcomes can be usefully represented by a network (Delmas et al., 2019; Newman, 2003). These networks convey information about which species achieve success in their interactions with other species. They can be represented as unipartite networks that characterise a single type of outcome, e.g. who eats whom in a food web; or bipartite networks, where interactions only occur between two distinct groups, such as plant–pollinator interactions. Estimating the minimum dimensionality of such interaction networks before deciding how many traits to investigate can prevent the omission of important traits. More accurate prediction of interaction outcomes could be made by combining information on the minimal number of necessary traits with methods to investigate the contribution of specific traits (Pichler et al, 2019). Minimum dimensionality can also inform theoretical network models about the minimum number of trait axes which have to be assumed for the reproduction of realistic uni- and bipartite ecological networks, ideally at the interaction outcome level (Olito & Fox, 2015).

The way traits participate in the interaction appears to be important for the estimation of minimum dimensionality, and we identify three relevant features. First, in a specific type of interaction, species in the community can exploit resources via different strategies, possibly employing different sets of traits, hence influencing minimum dimensionality. For example, flowering plants use visual or olfactory signals to achieve pollination by animals (Schiestl & Johnson, 2013), viruses can infect bacteria via alternative molecular pathways (Meyer et al., 2012), filamentous fungi employ different combative mechanisms in fungal competition for space (Hiscox & Boddy, 2017), and zooplankton species exhibit a variety of feeding modes such as active predation and filter feeding (Kiørboe, 2011). We refer to these alternative strategies for achieving a successful interaction outcome as ‘interaction modes’. Second, to exploit a resource successfully via a given interaction mode, an exploiter may have to succeed in different ‘tasks’, potentially employing different traits. For instance, a predator has to succeed in all the tasks of a typical predation sequence: encounter, detect, identify, approach, subjugate and consume a prey (Endler, 1991). Third, failure to exploit a resource is a trait-mediated outcome of an interaction, i.e. a ‘forbidden link’ or ‘forbidden interaction’ (Jordano, Bascompte, & Olesen, 2003; Morales-Castilla, Matias, Gravel, & Araújo, 2015). In this sense, two species interact given their mere inclusion in the network, even if they never actually meet, e.g. through temporal mismatch. Thus, traits involved in failures can also be included, which may differ from the traits involved in successes. Including these three features (alternate interaction modes, the number of tasks per mode and the trait-mediated failures) is expected to increase the number of traits involved in interactions, consequently increasing minimum dimensionality estimates.

Existing approaches for estimating minimum dimensionality lack at least one of these three features. To overcome these limitations, approaches should include a mechanistic description of interactions broken down to tasks, an explicit incorporation of the widespread feature of alternative interaction modes, or the consideration of failures as trait-mediated outcomes of interaction. Eklöf et al. (2013) developed a method for estimating minimum dimensionality in different interaction types, rooted in the theory of food web intervality in niche space (Cohen, 1977). In this method, dimensions originate phenomenologically, each dimension potentially accounting for multiple tasks and, hence, traits. Dalla Riva and Stouffer’s (2016) minimum dimensionality method builds on this approach by adopting a simple trait space representation for trophic interactions. Interaction network structure is explained mechanistically here by explicitly modelling exploiter–resource trait pairing and comparison. However, Dalla Riva and Stouffer (2016) model interactions via a single interaction mode since the task outcomes act additively from each corresponding exploiter– resource trait pair comparison. The same focus on a single interaction mode appears in Eklöf et al. (2013), because niche dimensions act in conjunction to determine the niche of the exploiter (resource). Lastly, it is common for behavioural studies to employ predictor traits to explain only the observed dominance events in a system, i.e. only the success outcomes (Chase & Seitz, 2011). Such studies commonly interpret interaction outcomes qualitatively, instead of quantifying minimum dimensionality, yet they can lead to incomplete explanations by overlooking other traits which might contribute only to the interaction failure outcomes.

Our aim is to develop a novel method which combines alternate interaction modes broken into mechanistically-based tasks and trait-mediated failure in a minimum dimensionality measure. Our ‘minimum mechanistic dimensionality’ measure can be applied to a broad range of uni- and bipartite ecological systems, including animal dominance, predator–prey, primary consumption, pollination, parasitism and seed dispersal networks, within a simple phenotypic trait space representation. We ask how our minimum dimensionality estimate compares to previous approaches across a range of different empirical ecological network types: under the assumption of alternative interaction modes compared to a single mode; with failure outcomes taken into account instead of omitted; and under a mechanistic perspective compared to the minimum dimensionality under the phenomenological, niche approach of Eklöf et al. (2013). Therefore, we test for potential underestimation of minimum dimensionality by previous approaches leading to the omission of relevant traits and mechanisms underlying interactions and community structure.

## 2 METHODS

Here, we illustrate our general approach with an empirical example of cyclic spatial replacement between three competing marine invertebrates. While the minimum dimensionality of this small, intransitive network is equal to one following Eklöf et al.’s (2013) methods, the estimate from our method equals two, providing a good example to illustrate our approach. We describe the interactions among species in the context of exploiter and resource roles, before going on to define and calculate the minimum mechanistic dimensionality of the network using a linear inequalities formulation. We then describe how we compared competing minimum dimensionality estimates across 658 empirical examples, ranging amongst social hierarchies, mutualistic networks and food webs.

### 2.1 Minimum mechanistic dimensionality: an overview of the method

Jackson and Buss (1975) described the cyclic spatial replacement of three marine, encrusting invertebrates: an ectoproct species *Stylopoma spongites* (player *A*) replacing a sponge species *Tenaciella* sp. (player *B*), which in turn replaces sponge species *Toxemna* sp. (player *C*), and which in turn replaces the ectoproct species player *A*. In our framework, a species can adopt the role of an exploiter, a resource, or exploiter-and-resource. In the marine invertebrates example, we consider any species both exploiter-and-resource of the other species, representing the observed replacement outcomes of spatial competition with a unipartite network (Fig. 1). Exploiters possess traits which participate in achieving exploitation, whereas resources possess traits working against resource exploitation. For success in a task, an exploiter’s performance in a given trait, termed ‘power’, must be higher than the resource’s performance in a corresponding trait called ‘toughness’ (taken from the creature combat rules of the card game *Magic: The Gathering*^®^ in Garfield, 2017). We consider that exploiter and resource are challenged in one trait ‘dimension’ of their phenotype space, where the performance of the corresponding power–toughness traits is directly compared, to determine who succeeds in that task. Using Boolean logic terms, interaction modes can be represented as OR-associated clauses of AND-associated tasks (see examples of one 2-dimensional mode and two 1-dimensional modes in Fig.1). Any structure of logical statements can be equivalently expressed in this ‘disjunctive normal form’ (Cohn, 2003), which we term the ‘interaction form’ here, providing a systematic description of how interactions occur.

**FIGURE 1.**
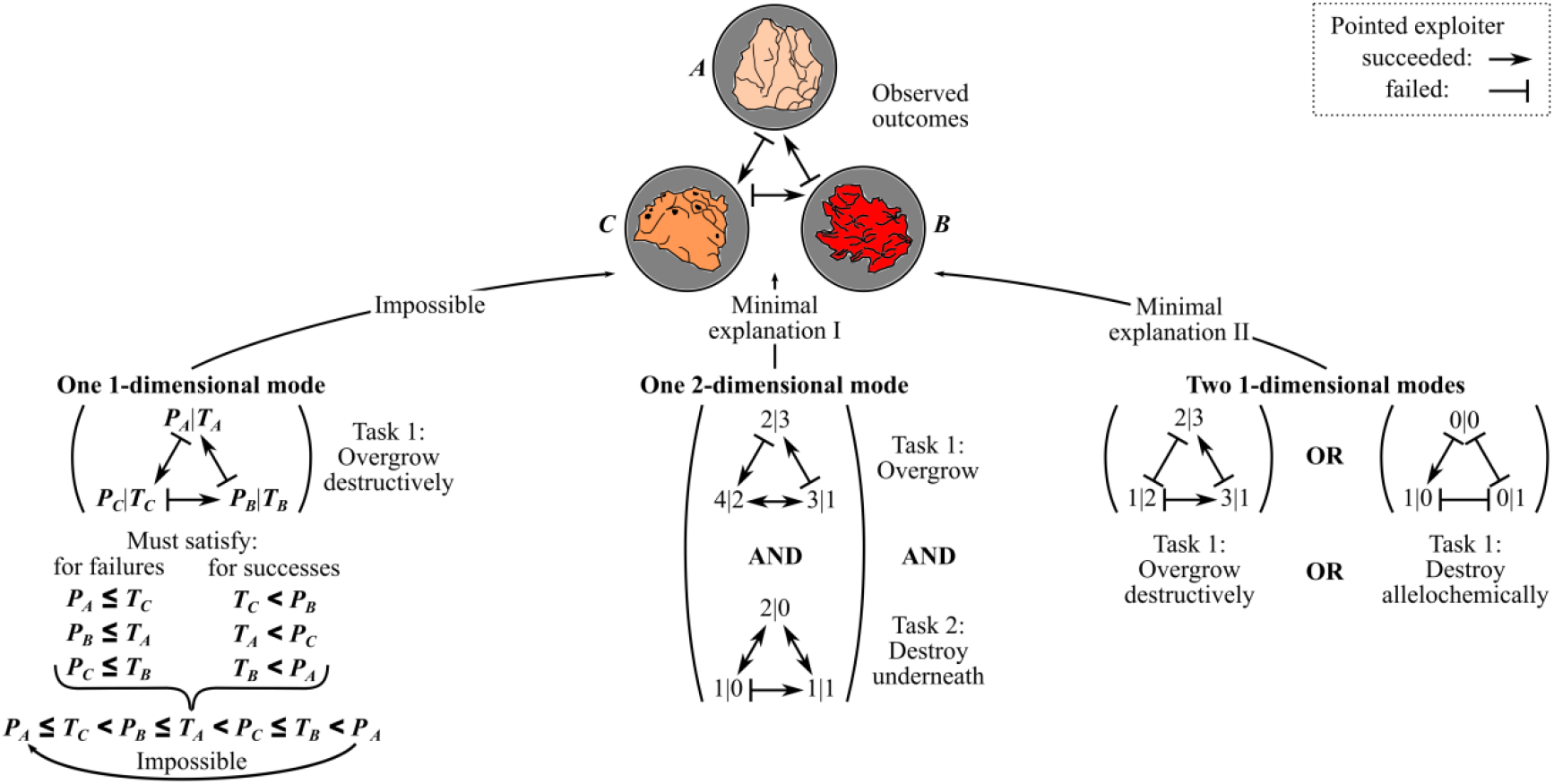
Explaining the observed outcomes of competition in an empirical rock–paper–scissors system of spatial replacement in three marine invertebrates. Each species was considered exploiter-and-resource of the others, hence possessing a power-and-toughness trait pair per task. We illustrate three attempts to minimally explain the observed outcomes: the first attempt with one 1-dimensional mode is mechanistically impossible, presuming a single trait pair for a single task, i.e. one dimension; the other two attempts are feasible, requiring two trait pairs in two tasks, i.e. two dimensions. We indicate hypothetical tasks, and power–toughness trait scores in arbitrary units of performance.

The only input for our method is the set of observed interaction outcomes (i.e., the observed spatio-temporal coexistence of the three species). We then define an interaction form describing the number of interaction modes which produced these outcomes, with each mode having a specific number of tasks. Since our aim is a minimum dimensionality estimate, we start with the simplest interaction form of a single task. In our example, we assume that interactions occurred via the destructive overgrowth of a rival invertebrate. For this task, a single pair of opposed, exploiter–resource power–toughness traits is assumed for all species. For example, the body height of the invertebrates when extending to an adjacent rival could be a trait for the power to overgrow destructively; and their body height when defending against overgrowth by rivals could be a trait for the toughness against destructive overgrowth. We then confront this trait pair in a system of inequalities, to satisfy the observed task successes and failures which coincide with the observed outcomes for this interaction form of a single task. For the task successes, the power of a winning exploiter must be greater than the toughness of a defeated resource, e.g. the exploiter’s body height must be higher than the defender’s body height. For the task failures, the power of a losing exploiter must be less than or equal to the toughness of an undefeated resource. In our example, the resulting system of six linear inequalities creates a cyclic sequence of ever-increasing power–toughness scores (the impossible ‘one 1-dimensional mode’ in Fig. 1). Thus, it is impossible to explain the observed outcomes in this unipartite graph if we presume that interactions occurred via a 1-dimensional interaction mode of a single task.

Our framework provides two alternative minimal mechanistic explanations for the emergence of this rock–paper–scissors network. First, we can find feasible power–toughness scores if we add a second task, i.e. another pair of power–toughness traits in the same mode (minimal explanation I in Fig. 1). We explain the failure of players *A* and *B* as failure in the first task (e.g. overgrowth), and the failure of *C* as failure in the second task (e.g. destruction of rival, even if *C* can overgrow *B*). Second, we can find solutions if we add a second interaction mode with one task, i.e. another pair of power–toughness traits in a new 1-dimensional mode (minimal explanation II in Fig. 1). In that case, we explain the success of *A* and *B* as success via the first mode (e.g. destructive overgrowth), and the success of *C* as success via a second mode (e.g. allelochemical elimination). Minimal explanation II is described by Jackson and Buss (1975) for this cryptic reef system: player *A* replaces *B* via overgrowth, player *B* replaces *C* via overgrowth, but player *A* is replaced by *C* due to toxic effects. Since the addition of a second task, i.e. power–toughness trait pair, leads to feasible power–toughness scores under both minimal explanations, the minimum mechanistic dimensionality of the empirical network equals two in both cases. Appendix S1 in the Supporting Information includes the complete systems of linear inequalities for the rock– paper–scissors network under minimal explanations I and II, formulated according to the method details presented next.

### 2.2 Minimum mechanistic dimensionality: formulating the inequalities

As illustrated with the rock–paper–scissors example (Fig. 1), the mechanistic explanation of the interaction outcomes in a network might require multiple tasks distributed across multiple interaction modes, i.e. requiring more than one pair of opposed exploiter– resource trait dimensions. One method to find this minimum number of trait dimensions is by attempting to solve a system of linear inequalities. If the system of linear inequalities is impossible, a simple strategy is to increase the number of dimensions (*d*) by one, and retry. Our minimum mechanistic dimensionality estimate is, therefore, the minimum *d* ≥ 1 for a feasible system of inequalities. In the marine invertebrates example of Fig. 1, there were two types of minimal explanation: additional trait pairs belonging to the same mode (minimal explanation I); or belonging to other, independent, 1-dimensional modes (minimal explanation II). We will focus on illustrating these two extreme explanations, although tasks could be distributed to interaction modes in various other ways for cases requiring more than two tasks.

When a new task is added to a single mode, permitting feasibility of the system of linear equalities, the *d* exploiter–resource trait pairs (dimensions) must be involved in the same mode (minimal explanation I, Fig. 1). On one hand, an observed success of exploiter *A* against resource *B* must be the result of success in all tasks (e.g. player *A* succeeds in both overgrowing and destroying *B*). Specifically, the power *P*_*A,i*_ ≥ 0 of exploiter *A* in any trait pair *i* must be greater than the toughness *T*_*B,i*_ ≥ 0 of resource *B* in that trait pair: *P*_*A,i*_ > *T*_*B,i*_. Since success might require more than the marginal superiority of the exploiter’s power, we add a superiority threshold, *t*_*A,B,i*_ > 0, making the task success requirement:

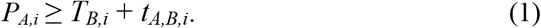

On the other hand, the observed failure of exploiter *A* against resource *B* must be the result of failure in at least one task (e.g. player *C* failing at task 2 against *B* in Fig. 1). We can use a binary variable as an indicator of failure in trait pair *i*, *f*_*A,B,i*_ (Williams, 2013). If *f*_*A,B,i*_ = 1, then exploiter *A* fails against resource *B* in trait pair *i*; otherwise, *f*_*A,B,i*_ = 0, representing exploiter success in the task. Finally, we include bounds for the power–toughness differences for computational efficiency (Williams, 2013): a sufficiently negative lower bound *m* of the exploiter’s power inferiority in case of task failure; and a sufficiently positive upper bound *M*, of the exploiter’s power superiority in case of task success. Here, we set *m* = −200 and *M* = 200, but these limits were not reached in any of the empirical networks we considered below. Thus, for an observed failure, the following pair of inequalities must be satisfied in any trait pair *i*:

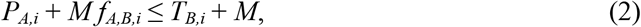

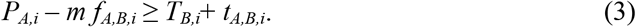

With an extra inequality for the observed failure, we can force at least one of the binary indicator variables to equal one, i.e. failure in at least one task:

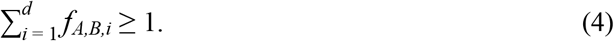

In the case of a task failure in trait pair *i*, *f*_*A,B,i*_ = 1, inequality (2) is the task failure requirement, and inequality (3) is the lower bound for the exploiter’s power inferiority. In case of a task success, *f*_*A,B,i*_ = 0, inequality (2) gives the upper bound for the exploiter’s power superiority, and inequality (3) becomes a success requirement.

When a new 1-dimensional mode is added, permitting a feasible system of linear inequalities (minimal explanation II, Fig. 1), each of the *d* pairs of opposed exploiter–resource traits must be involved in a different mode. On one hand, the observed failure of any exploiter *A* against any resource *B* must be the result of failure in any mode *i* of the *d* modes (e.g. player *A* failing via both overgrowth and allelopathy against *C* in Fig. 1):

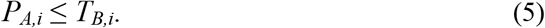

On the other hand, the observed success of exploiter *A* against resource *B* must come from success via at least one mode (e.g. player *C* replacing *A* via allelopathy in Fig. 1). We now use a binary variable, *s*_*A,B,i*_, to indicate success via mode *i*. Given the same bounds as in minimal explanation I, the following pair of inequalities must be satisfied to indicate exploiter success in any mode *i*:

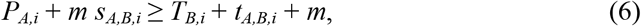

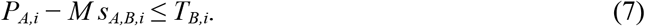

With an extra inequality for the observed success, we can force at least one of the binary indicator variables to equal one, i.e. exploiter success occurs via at least one interaction mode:

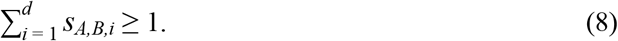

A complete system of linear inequalities takes into account all observed successes and failures for all possible exploiter–resource pairs. Such systems of linear inequalities, with continuous trait values and integer indicator variables, can be formulated and attempted to be solved as mixed integer linear programming problems (Williams, 2013). In both minimal explanations (I and II), minimum mechanistic dimensionality is the minimum *d* leading to a feasible system of inequalities (see Fig. 2 for a summary of the method’s procedure in pseudocode).

**FIGURE 2.**
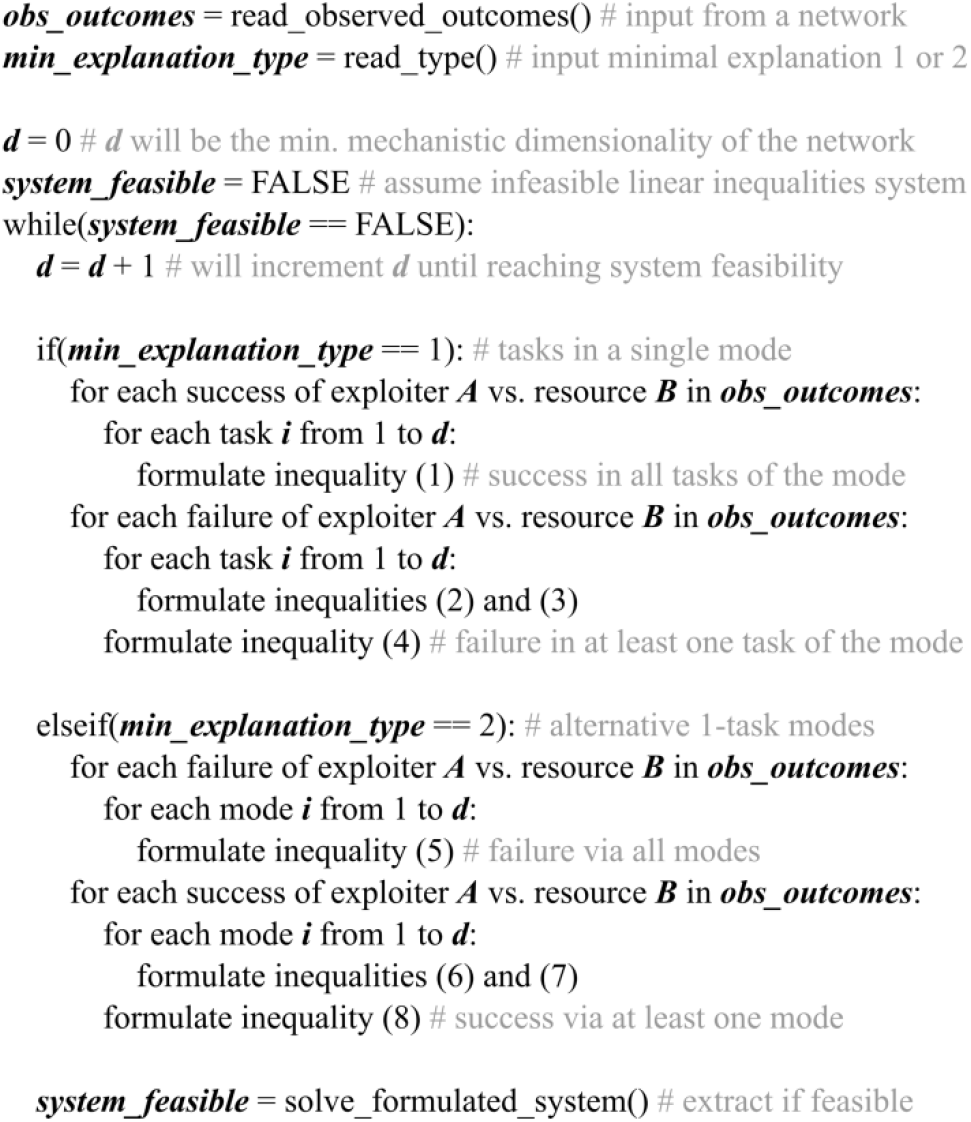
Pseudocode for estimating the minimum mechanistic dimensionality of an ecological network from the observed outcomes under minimal explanations I and II.

### 2.3 Comparing the minimum dimensionality of empirical networks

We applied our method to 658 empirical systems, covering six different types of ecological networks: animal social dominance networks, food webs with basal species excluded, basal–consumer interactions, plant–pollinator, host–parasite and seed dispersal networks (see Appendix S1 for details). Considering a single interaction type in each system, and assuming adequate sampling effort (e.g. no observed failures due to rarity), we computed five dimensionality measures (Appendix S1). Four minimum mechanistic dimensionality measures were based on our framework: (*a*) a single, potentially multidimensional mode; (*b*) one-or-more 1-dimensional modes; (*c*) as measure *b*, but excluding observed failures; (*d*) as measure *c*, but with players interacting via a common trait per dimension, rather than a power against toughness trait. To compare our approach with another established dimensionality estimate in this first account, the fifth measure we considered (*e*) was that of Eklöf et al. (2013). We used these five dimensionality measures to answer three questions about minimum mechanistic dimensionality (MMD): (1) Is MMD higher under the assumption of alternative 1-dimensional modes (dimensionality measure *b*), compared to the assumption of a single multidimensional mode (dimensionality measure *a*)? (2) Is MMD higher with observed failures taken into account (dimensionality *b*), or excluded (dimensionality *c* or *d*)? (3) Is MMD higher than the dimensionality measure developed by Eklöf et al. (2013) (dimensionality *a* versus *e*)?

The systems of linear inequalities for our four minimum mechanistic dimensionality measures *a–d* were formulated and solved as mixed integer linear programming problems with the Gurobi Optimizer (Gurobi Optimization and Inc., 2017). R and Python code for formulating and solving for these four minimum mechanistic dimensionalities are provided (see section ‘DATA ACCESSIBILITY’). We computed the fifth dimensionality *e* with code provided in the Supporting Information of Eklöf et al. (2013). The references we provide for the empirical networks were retrieved from five data sources (Cohen, 2010; Ortega, Fortuna, & Bascompte, 2017; Shizuka & McDonald, 2015; Stanko & Miklisova, 2014; Thompson & Townsend, 2004). We provide network characteristics and references, and raw data from the five computed dimensionality measures, for each of the 658 empirical systems (see section ‘DATA ACCESSIBILITY’).

## 3 RESULTS

For the five dimensionality measures we considered, the inclusion of alternative interaction modes, forbidden links and a mechanistic approach to describing interactions consistently increased the minimum dimensionality estimate across a wide range of empirical ecological networks.

### 3.1 Unimodal *vs* multimodal minimum mechanistic dimensionality

Our estimate of minimum mechanistic dimensionality was frequently higher under the alternative modes explanation than under the single mode explanation (Fig. 3), especially in systems of non-basal consumption, biotic pollination, ectoparasitism, and seed dispersal (Fig. 3b,d–f). 54% of the empirical systems had higher dimensionality if alternative modes were assumed, with only 7% of the systems having higher unimodal dimensionality (Fig. 4a).

**FIGURE 3.**
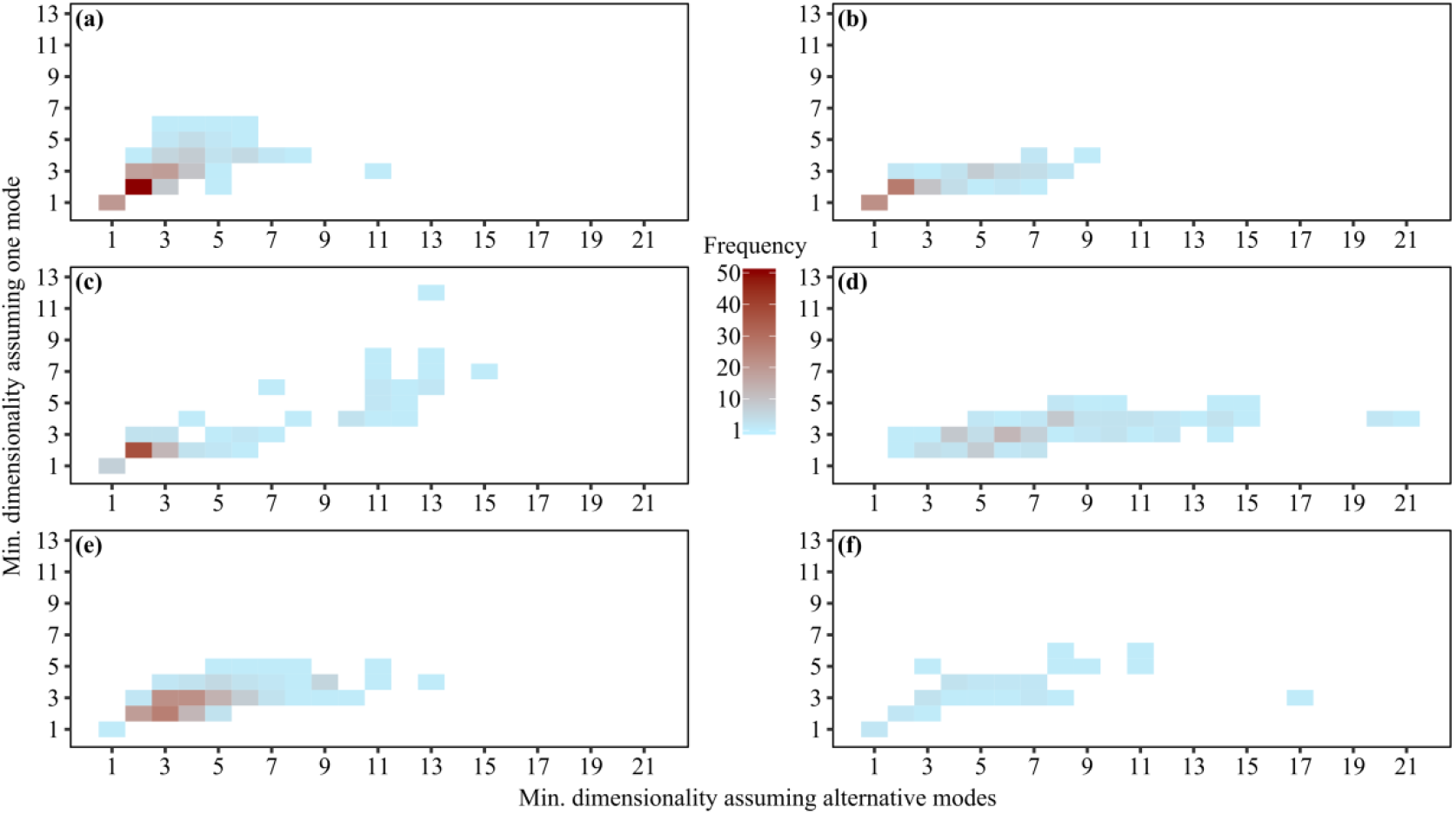
Minimum mechanistic dimensionality estimates from 658 empirical systems. Cell shading indicates how many of the *n* systems had the corresponding pair of values in our two minimum dimensionality measures, i.e. number of exploiter–resource trait pairs pre-assuming: alternative 1-dimensional modes (x-axis; minimal explanation II); tasks in a single mode (y-axis; minimal explanation I). We focused on one interaction type in each empirical system: (a) animal dominance in *n* = 168 unipartite graphs (of 6–31 individuals); (b) consumption of non-basal species in *n* = 95 unipartite food webs (6–57 species, basal species excluded from the original food webs); (c) consumption of basal species by consumers exclusively feeding on them in *n* = 95 bipartite graphs (11– 91 species; from the same food webs used in (b)); (d) plant biotic pollination in *n* = 105 bipartite graphs (8–114 species); (e) ectoparasitism of small mammals in *n* = 165 bipartite graphs (8–92 species); and (f) plant seed dispersal in *n* = 30 bipartite graphs (6–86 species). Parameter values in the linear inequalities method: *m* = −200; *M* = 200; *t*_*A,B,i*_ = 1, for all pairs of exploiter *A* with resource *B*, and in any trait pair *i*.

**FIGURE 4.**
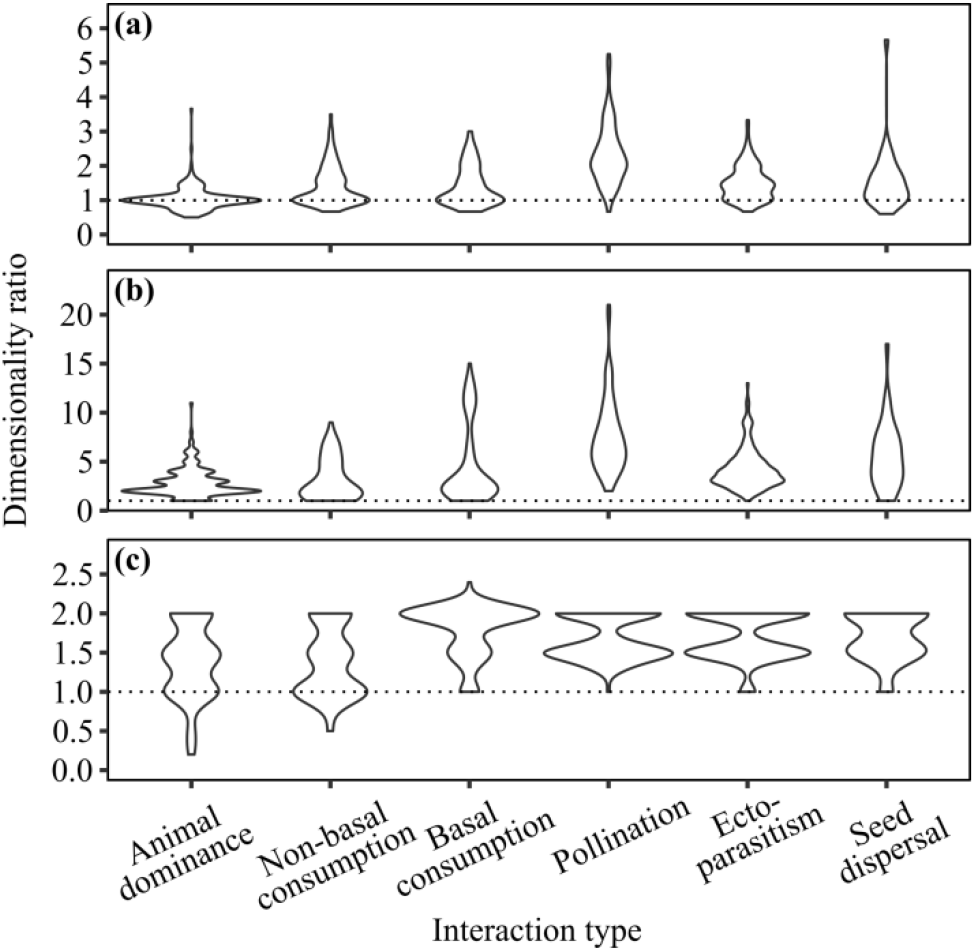
Comparisons of minimum dimensionality measures estimated from 658 empirical systems. For each empirical network, we calculated the ratio of: (a) our minimum mechanistic dimensionality under minimal explanation II (alternative 1-dimensional modes), to our minimum mechanistic dimensionality under minimal explanation I (tasks in a single mode); (b) our minimum mechanistic dimensionality under minimal explanation II, to the same dimensionality with the failures ignored by the system of linear inequalities; and (c) our minimum mechanistic dimensionality under minimal explanation I, to the comparable dimensionality of Eklöf et al. (2013). Violin plots show the normalised distributions of the dimensionality ratios. Dotted horizontal lines show a ratio of one.

### 3.2 Minimum mechanistic dimensionality with *vs* without failure outcomes

Minimum mechanistic dimensionality was higher in 92% of the empirical systems when failure outcomes were included, instead of excluded (Fig. 4b). In the remaining 8% of empirical systems, the two dimensionality estimates were equal. Here, we compared our minimum multimodal dimensionality with the same dimensionality but with any failure inequalities excluded from the inequalities system. When failures were excluded, the minimum dimensionality always equaled one. In this case, the structure of observed successes can be explained unimodally, as exploiters can have a single power trait with a greater value than the single toughness trait of any resource (in the absence of any inequalities constraining the power scores). We further required that exploiters and resources possess the same trait for power and toughness in the unipartite systems of animal dominance and non-basal consumption, instead of the default power–toughness trait pair. In that way, the unipartite systems could require more than one dimension with failures excluded. Even in this case of modelling trait opposition with a single, common trait per dimension, 79% of the unipartite systems had higher minimum dimensionality with failures included rather than excluded.

### 3.3 Mechanistic *vs* phenomenological minimum dimensionality

In 81% of the empirical systems, our minimum mechanistic dimensionality was higher than the dimensionality estimate of Eklöf et al. (2013). We assumed a single mode (minimal explanation I), which is comparable to the niche approach of Eklöf et al. (2013). Only 2% of the networks had higher minimum dimensionality under the more phenomenological approach of Eklöf et al. (2013), with no bipartite networks among them (Fig. 4c).The minimum number of trait pairs for the explanation of all outcomes in our approach, was 1.5 times larger (median) than with the phenomenological dimensionality across all networks. Note that since our dimensionality refers to exploiter–resource trait pairs, the actual number of necessary traits is double our dimensionality, i.e. the present approach suggested a median of 3 times more trait axes required for the explanation of the observed outcomes than the Eklöf et al. (2013) approach.

## 4 DISCUSSION

We introduced a method for calculating the minimum number of traits required to explain all observed interaction outcomes of ecological networks mechanistically, using a general framework applicable to different interaction types, modes, tasks, and types of traits. Applying this to 658 empirical systems, we showed that the minimum number of traits involved is typically underestimated when ignoring any of the three framework features we combined here for the first time: (1) alternative interaction modes; (2) trait-mediated failure outcomes; and (3) a mechanistic description of interactions. This underestimation risks omitting important traits in empirical investigations, and generating less realistic theoretical networks at the interaction outcome level.

With our generalising framework, minimum mechanistic dimensionality can explicitly incorporate the alternative interaction modes frequently observed empirically, e.g. alternative feeding modes. In previous theoretical trait-based works, an exploiter has to overcome all the barriers or defences of a potential resource to consume or parasitise the resource (Gilman, Nuismer, & Jhwueng, 2012; Santamaría & Rodríguez-Gironés, 2007). Similarly, in other theoretical works adopting a niche approach, a niche arises from the intersection of all niche dimension intervals (Eklöf et al., 2013; Stouffer, Camacho, & Amaral, 2006). The interaction mode in our framework is equivalent to these two approaches, as an exploiter’s performance must be sufficiently high in all the mode’s tasks. Additionally, by generalising to alternative interaction modes, we showed that minimum mechanistic dimensionality can differ, frequently being higher under alternative interaction modes than in a single mode (Fig. 4a). Moreover, different assumptions about the interaction form can also lead to alternative minimal explanations of the outcomes, as in rock–paper–scissors systems (Fig. 1), offering a new mechanistic perspective in the study of intransitive networks (Szolnoki et al., 2014).

We adopted a phenotype rather than a niche space representation for traits. Studies of interactions commonly use the ‘resource-utilization’ approach to represent the ‘ecological niche’ concept (MacArthur & Levins, 1967, Schoener, 1989). Despite its operational advantage (Schoener, 1989), dimensions usually arise phenomenologically, as in the minimum dimensionality of Eklöf et al. (2013). For example, body size is a trait with high explanatory power in food webs (Stouffer, Rezende, & Amaral, 2011). However, more traits that scale allometrically with body size are mechanistically involved in trophic interactions (Woodward et al., 2005). Even if taken mechanistically, realised niches commonly span a range of the resource gradient (MacArthur & Levins, 1967), implying two traits per niche dimension. For instance, in systems where prey size range is limited by a predator’s mouth gape, the niche range minimum must be limited by a second trait, like the predator’s inability to handle smaller prey. Another problem with the resource-utilization approach is that exploiters are implicitly excluded from the niche space, as it is created by trait dimensions of the resources (MacArthur & Levins, 1967; Schoener, 1989). Our framework accounts for the traits of both interacting players simultaneously, and a dimension is simply a challenged trait-axis in the phenotype space of exploiters and resources, as in the approach of Dalla Riva and Stouffer (2016). In that way, we found that our minimum mechanistic dimensionality assuming a single interaction mode was frequently higher than the comparable dimensionality of Eklöf et al’s (2013) niche-based approach (Fig. 4c).

We regarded failures as trait-mediated outcomes of an interaction, meaning more traits were expected to be involved in the interactions (Fig. 4b). We found that three to six pairs of opposed traits must be involved in several behavioural dominance systems (Fig. 3a), whereas only a few traits are commonly employed for the explanation of the successful dominance outcomes (Chase & Seitz, 2011). For example, in the elephant family named ‘AA’ in Archie et al. (2006), almost all observed dominance events were directed towards younger elephants, and the authors conclude the system is an age-ordered dominance hierarchy based only on the successes, in agreement with our one dimension with failures-excluded analysis (Fig. 4b). However, our minimum mechanistic dimensionality, explicitly incorporating failures, suggests three trait pairs under both minimal explanations, because there are several older– younger pairs where no dominance or aggression was observed, i.e. failures unaccounted for by Archie et al. (2006). In fact, most elephants dominated only younger members within their matriline, and within two specific matrilines (Archie et al., 2006). These two characteristics are candidates for the two extra dimensions predicted by our method, which are overlooked when ignoring failure outcomes.

In this first account, we assumed two simple and extreme minimal interaction forms (minimal explanations I and II), but the user can input any minimum number of traits and trait values, in any interaction form. Additionally, we presented a deterministic version, but future versions could be probabilistic (Dalla Riva & Stouffer, 2016; Poisot et al., 2016), e.g. with more probable outcomes explained by larger power–toughness differences. Assuming adequate sampling effort, our mechanistic description has not considered the effects of abundance (Vázquez & Aizen, 2003), which could also be incorporated in future extensions. Lastly, our method assumes that performance is independent in the different tasks, i.e. a unique performance trait per task per player in our formulation of the linear inequalities. In reality, several traits of an individual can contribute to performance in the same task, and the same trait can contribute to performance in several tasks (Arnold, 1983). Since our aim was a minimum dimensionality measure, we assumed independence in task performance to impose fewer constraints in the linear inequalities system, allowing the estimation of a lower minimum.

In conclusion, we have outlined a novel method of estimating the minimum dimensionality of ecological networks, which is relatively straightforward to adopt and calculate. Informed by a more accurate minimum dimensionality estimate, future studies can rely on network models reproducing community structure more accurately at the interaction outcome level (Olito & Fox, 2015), and can reduce the risk of missing important traits that are involved mechanistically, in alternative interaction modes, or only in failure outcomes. In that way, our method could contribute to a better understanding, explanation, and prediction of community structure and structure-dependent processes.

## ACKNOWLEDGEMENTS

The authors would like to thank David Gilljam and Miguel Lurgi Rivera for comments which improved the manuscript. Swansea University supported DAK with a student scholarship.

## AUTHORS’ CONTRIBUTIONS

DAK developed the method and the required code, compiled the empirical networks dataset, computed the dimensionalities of the networks, analysed the data, and wrote the first draft of the manuscript. All authors contributed to the development of the method, data analysis, and to the writing of the manuscript.

## DATA ACCESSIBILITY

Data and code will be made available at the Dryad Digital Repository.

